# Rice blast resistance gene *Pii* is controlled by a pair of NBS-LRR genes *Pii-1* and *Pii-2*

**DOI:** 10.1101/227132

**Authors:** Hiroki Takagi, Akira Abe, Aiko Uemura, Kaori Oikawa, Hiroe Utsushi, Hiroki Yaegashi, Hideko Kikuchi, Motoki Shimizu, Yoshiko Abe, Hiroyuki Kanzaki, Hiromasa Saito, Ryohei Terauchi, Koki Fujisaki

## Abstract

Nucleotide-binding, leucine-rich repeat receptors (NLRs) are conserved cytosolic receptors that recognize pathogen effectors and trigger immunity in plants. Recent studies indicate that NLRs function in pairs. Rice resistance gene *Pii* has been known to confer resistance against rice blast pathogen *Magnaporthe oryzae* carrying *AVR-Pii*. Previously we reported isolation of *Pii* gene from the rice cultivar Hitomebore (Takagi et al. 2013). To further understand rice components required for *Pii*-mediated resistance, we screened 5,600 mutant lines of Hitomebore cultivar and identified two mutants that lost *Pii* resistance without any changes in *Pii* gene sequence. Application of MutMap-Gap, the whole genome sequencing-based method of mutation identification, to the two mutants revealed that they have mutations in another *NLR* gene located close to *Pii*. The F1 plants derived from a cross of the two mutants showed *pii* phenotype, demonstrating that the newly identified *NLR* gene is indeed a component of *Pii* resistance. We thus designate the previously isolated *Pii* gene as *Pii-1* and the newly isolated NLR gene as *Pii-2*.

## INTRODUCTION

Plants are facing attacks from various pathogenic microbes in natural environments. Pathogens secrete a battery of virulence effectors to manipulate host function. As a counter-strategy, plants evolved resistance (*R*) genes whose products recognize pathogen avirulence effectors (AVRs). The recognition of AVRs leads to a strong defense response known as effector-triggered immunity (ETI). In most cases R proteins belong to the protein family of nucleotide-binding, leucine-rich repeat receptors (NLR) that directly or indirectly recognize AVRs (Win *et al*., 2012).

Mechanistic function of NLRs has not been fully understood. However, an increasing number of evidences show that NLRs cooperate with genetically linked or unlinked NLRs (Duxbury *et al*., 2016; Wu *et al*. 2017). For example, *RPS4/RPS1, RGA4/RGA5* and *Pikp-1/Pikp-2* are genetically linked *NLR* gene pairs and both of the paired NLRs are required for the recognition of cognate AVRs. One of the NLR pair is called “sensor” that recognize AVRs and the other NLR is called “helper” that is needed for the immune signaling (Duxbury *et* al., 2016). Genetically unlinked pairs have been also identified. Solanaceae R-genes *Rbi-blb2, Mi-1,2* and *R1* require *NRC4 NLR* gene in an unlinked locus and *Prf* R-gene requires *NRC2* or *NRC3 NLRs* (Wu et al. 2017), leading to the proposa of the concept of “NLR network” (Wu et al. 2017).

Rice blast caused by the fungal pathogen, *Magnaporthe oryzae*, is the most destructive disease of rice (Dean *et al*., 2012; Zeigler *et al*., 1994) and the use of *R* genes in rice breeding is the most cost-effective measure to fight the disease (Ishizaki *et al*., 2005; Miah *et al*., 2013).

Among the cloned rice R-genes against *M. oryzae, Pia* and *Pik* have been reported to be composed of paired *NLRs* (Ashikawa *et* al., 2009; Okuyama *et* al., 2011; Yuan *et* al., 2011; Zhai *et al*., 2011). Their cognate AVRs, *AVR-Pia* and *AVR-Pik*, have also been isolated from *M. oryzae* (Yoshida *et al*., 2009). *Pia* consists of a pair of genetically linked *NLR* genes, *RGA4* and *RGA5*. RGA4 induces AVR-Pia-independent cell death, which is suppressed by RGA5. RGA5 directly interacts with AVR-Pia, and the RGA5-AVR-Pia interaction relieves the suppression of RGA4 function by RGA5 (Cesari *et al*., 2013). In this model, RGA5 functions as cell death suppressor and AVR sensor, whereas RGA4 functions as executor of defense responses. However, we still do not know whether this model proposed for *Pia* applies to other linked paired NLRs.

We have been studying *Pii*-mediated resistance of rice-*M. oryzae* interaction. In contrast to direct AVR recognition in *Pia-* and *Pik*-mediated resistance (Cesari *et al*., 2014; Kanzaki *et al*., 2012), indirect recognition of the AVR-Pii effector by Pii has been proposed (Fujisaki *et al*., 2015). By using MutMap-Gap, a whole genome sequencing-based method of mutation identification, we have isolated an *NLR* gene (*Pii*). In this study, we searched rice genes involved in *Pii*-mediated resistance, and identified another *NLR* gene required for *Pii*-mediated resistance. This new *NLR* gene as designated as *Pii-2* is different from the previously-identified *Pii* gene (Takagi *et al*., 2013), which is here renamed to *Pii-1*, but is genetically linked to the *Pii-1* locus.

## RESULTS AND DISCUSSION

### Screen of rice mutants showing deficiency of *Pii*-mediated resistance

The rice cultivar Hitomebore is known to have *Pii* gene and shows resistance to rice blast fungus carrying *AVR-Pii* (Yoshida *et al*., 2009). In the previous study, we isolated Hitomebore ethylmethanesulfonate (EMS)-induced mutants (Hit5948 and Hit6780) showing susceptibility to an incompatible isolate of *M. oryzae* with *AVR-Pii* (TH68-126). These mutants had loss-of-function mutations in the same *NLR* gene and we concluded it is *Pii* (Takagi *et* al., 2013). In the current study, to identify other genes required for Pii-mediated resistance, we newly screened a total of 5,600 Hitomebore EMS mutant lines by spray inoculation with TH68-126, resulting in the isolation of five mutant lines (Hit5882, Hit12904, Hit13537, Hit13701 and Hit14409) showing susceptibility to TH68-126. In these mutant lines, genomic sequence of the previously identified *Pii* gene (Takagi *et al*., 2013) was determined by Sanger sequencing. Three of the five mutant lines had mutations in the *Pii* gene (Hit12904: exon-intron junction; Hit13537: Q540stop; and Hit14409: exon-intron junction), suggesting that these mutations contributed to their susceptibility to TH68-126. However, other two mutants, Hit5882 and Hit13701, had no mutation in the *Pii* gene. The susceptible phenotypes of Hit5882 and Hit13701 to the incompatible isolate TH68-126 were confirmed by punch inoculation test (Fig. 1 and S1).

**Fig. 1.**
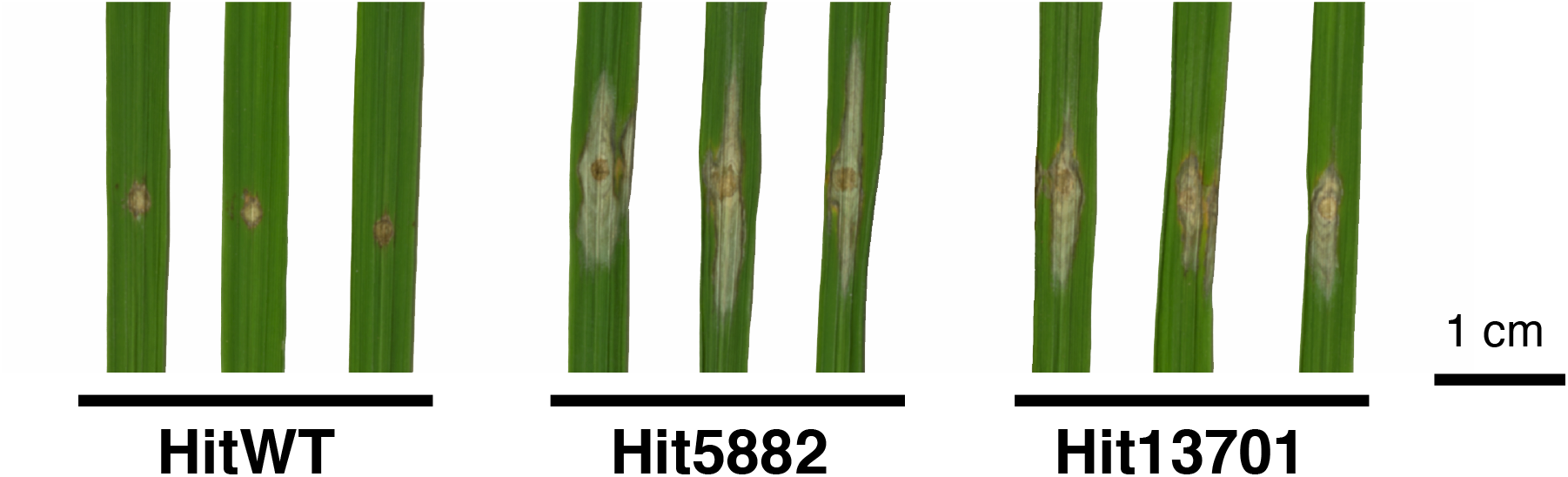
Rice Hitomebore mutants Hit5882 and Hit13701 deficient in *Pii*-mediated resistance. Rice leaf blade punch inoculation of *M. oryzae* isolate, TH68-126 (AVR-Pii+), to wild-type Hitomebore (HitWT: *Pii*+) and its mutants (Hit5882 and Hit13701). Inoculated leaves were photographed at 10 days after inoculation. HitWT shows resistance to TH68-126, whereas the two Hitomebore mutants, Hit5882 and Hit13701 are susceptible.

### Identification of *Pii-2* by MutMap-Gap

To identify the causative mutations of Hit5882 and Hit13701, we carried out MutMap-Gap (Fig. S2). First, the genomic regions harboring candidate mutations of the two mutants were identified by MutMap analysis (Abe *et al*., 2012). To apply MutMap, Hit5882 and Hit13701 were separately crossed with the wild type Hitomebore (HitWT) to generate the F2 progeny (hereafter designated as Hit5882 F2 family and Hit13701 F2 family, respectively). Phenotype of each F2 progeny was studied by punch inoculation with an incompatible *M. oryzae* isolate TH68-126 carrying *AVR-Pii*. The F2 progeny of Hit5882 family segregated to resistance (R) : susceptible (S) in the ratio of 86 : 33 and that of Hit13701 family segregated to R : S in the ratio of 83 : 22, both conforming to the 3 : 1 ratio (Chi square test; Hit5882 family: *χ*^2^ = 0.47, ns; Hit13701 family: *χ*^2^ = 0.92, ns), indicating that the susceptible phenotypes of both mutants were caused by single recessive mutations. For each family, DNA from all the susceptible F2 progeny (33 and 22 individuals for Hit5882 and Hit13701 family, respectively) were extracted and bulked in equal amount and subjected to whole genome sequencing by illumina NextSeq500 sequencer, resulting in 5.81 and 6.73 Gb sequence reads for Hit5882 and Hit13701 family, respectively (Table S1). The sequence reads were aligned by BWA software (Li *et al*., 2009) to “Hitomebore reference sequence”, developed by replacing the nucleotides of publically available Nipponbare reference genome (Os-Nipponbare-Reference-IRGSP-1.0) with those of Hitomebore at all SNP positions between the two cultivars. The SNP-index values (Abe *et al*., 2012), representing the allele frequency of mutant nucleotides among the F2 pool, were calculated at all SNP positions using MutMap pipeline ver 1.4.4 (http://genome-e.ibrc.or.jp/home/bioinformatics-team/mutmap). MutMap analysis of the two families showed single peaks in the genomic region from 9.5 to 13 Mb on Chromosome 9 (Fig. 2 and Fig. S3), which coincided with the previously-identified *Pii* gene (Takagi *et al*., 2013). None of the SNPs with SNP-index = 1 caused amino acid changes or corresponded to splicing junctions of the putative genes located in the region. Since the MutMap analysis was carried out on the “Hitomebore reference sequence” developed from the Nipponbare reference genome by nucleotide replacements, the SNPs present in Hitomebore-specific genomic region were not addressed in this analysis. Our failure of identification of candidate mutations in the region suggests that the causative mutation may be located on such Hitomebore-specific genomic region that are missing from the “Hitomebore reference sequence”.

**Fig. 2.**
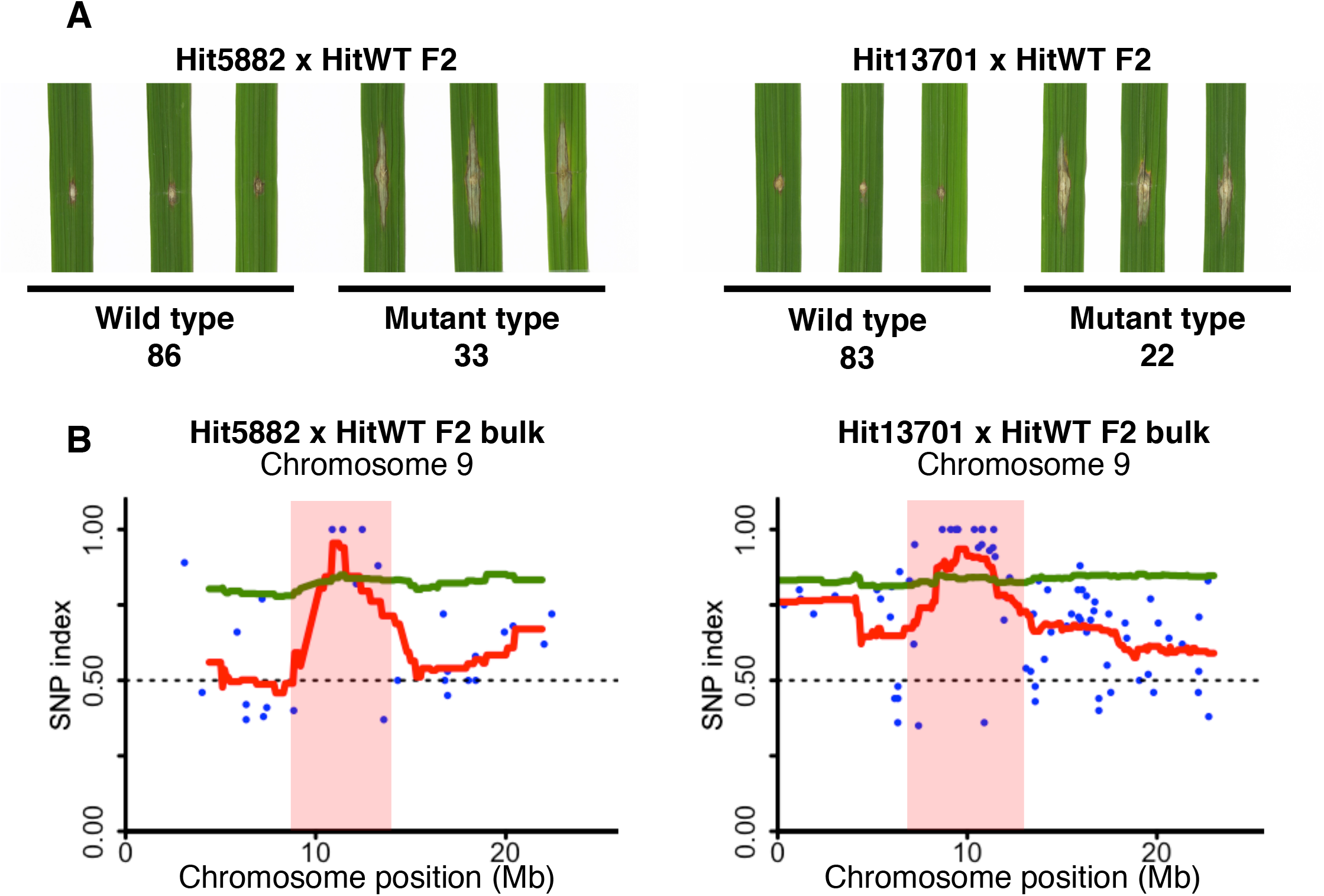
MutMap analysis of the mutant Hit5882 and Hit13701. (A) Segregation of wild-type and mutant phenotypes in the F2 families derived from a cross of the wild-type Hitomebore (HitWT) with Hit5882 (left) and that of HitWT with Hit13701 (right). Phenotype of F2 individuals was confirmed by punch inoculation of *M. oryzae* isolate TH68-126 carrying *AVR-Pii*. The numbers of F2 individuals showing wild-type (resistant) and mutant (susceptible) phenotypes are indicated at the bottom. (B) SNP-index plots of chromosome 9 in MutMap analysis for the mutations of Hit5882 and Hit13701. SNP-index for SNPs are given in blue dots. The red line indicates the sliding window average of SNP-index values of SNPs in 4 Mb interval with 10 kb increment. Green line represents the 95% statistical confidence limit under the null hypothesis of SNP-index = 0.5.

To search for the mutations localized in Hitomebore specific genomic regions, we carried out local *de novo* assembly to reconstruct Hitomebore specific sequences corresponding to the Nipponbare 9.5–13 Mb genomic region on Chromosome 9. To this end, the Hitomebore sequence reads mapped to 9.5–13 Mb on Chromosome 9 (total 168.54 Mb in size) and those unmapped to “Hitomebore reference sequence” (total 688.19 Mb in size) were extracted, and were together subjected to *de novo* assembly using DISCOVAR *de novo* (Weisenfeld *et al*., 2014). This *de novo* assembly resulted in the generation of 2,087 contigs amounting to the total size of 6,234 Kb with N50 value of 3,431 bp (Fig. S4). Combined with “Hitomebore reference sequence”, the sequences of the 2,087 contigs served as a new reference sequence. We aligned the bulked DNAs of mutant F2 of the Hit5882 family and Hit13701 family, respectively, to the new reference sequence.

MutMap-Gap analyses of Hit5882 and Hit13701 using the new reference sequence identified candidate mutations (SNP-index=1) in the same contig “no.298” that was generated by *de novo* assembly (Table S2), which harbored a sequence corresponding to a truncated gene encoding a putative NLR protein. Since the gene in contig “no.298” was partial, the 5’ and 3’ termini of the cDNA sequence were determined by Rapid Amplification of c-DNA Ends (RACE) method. Based on the sequence information of both the 5’-and 3’-UTR regions, the full-length protein coding region was amplified by PCR, and the sequence was determined by Sanger method (Fig. S5). Next, to determine the genomic sequence of the NLR gene, the obtained cDNA sequence was subjected to BLAST search using a database of the contigs developed by *de novo* assembly. The BLAST search revealed that contig 298 and contig 1164 showed similarity to the region 1-2,581bp and 3,256-3,717 bp of the *NLR* coding sequence, respectively, indicating that contig 298 and contig 1164 encode the upstream and downstream regions of the *NLR* genomic DNA, respectively. On the other hand, no contigs containing the internal region (2,582-3,255 bp) of the *NLR* coding sequence was recovered by BLAST. We therefore PCR amplified this region and sequenced by Sanger method. The resulting 5’-, 3’- and internal regions of the gene were assembled and finally 7,946 bp genomic DNA carrying the *NLR* gene of Hitomebore cultivar was determined (Fig. S5). The candidate mutation in Hit5882 causes an amino acid substitution (Val to Asp: V371D) and that in Hit13701 causes a nonsense mutation (Q247stop) in the NLR gene (Fig. 3 and Table S2).

**Fig. 3.**
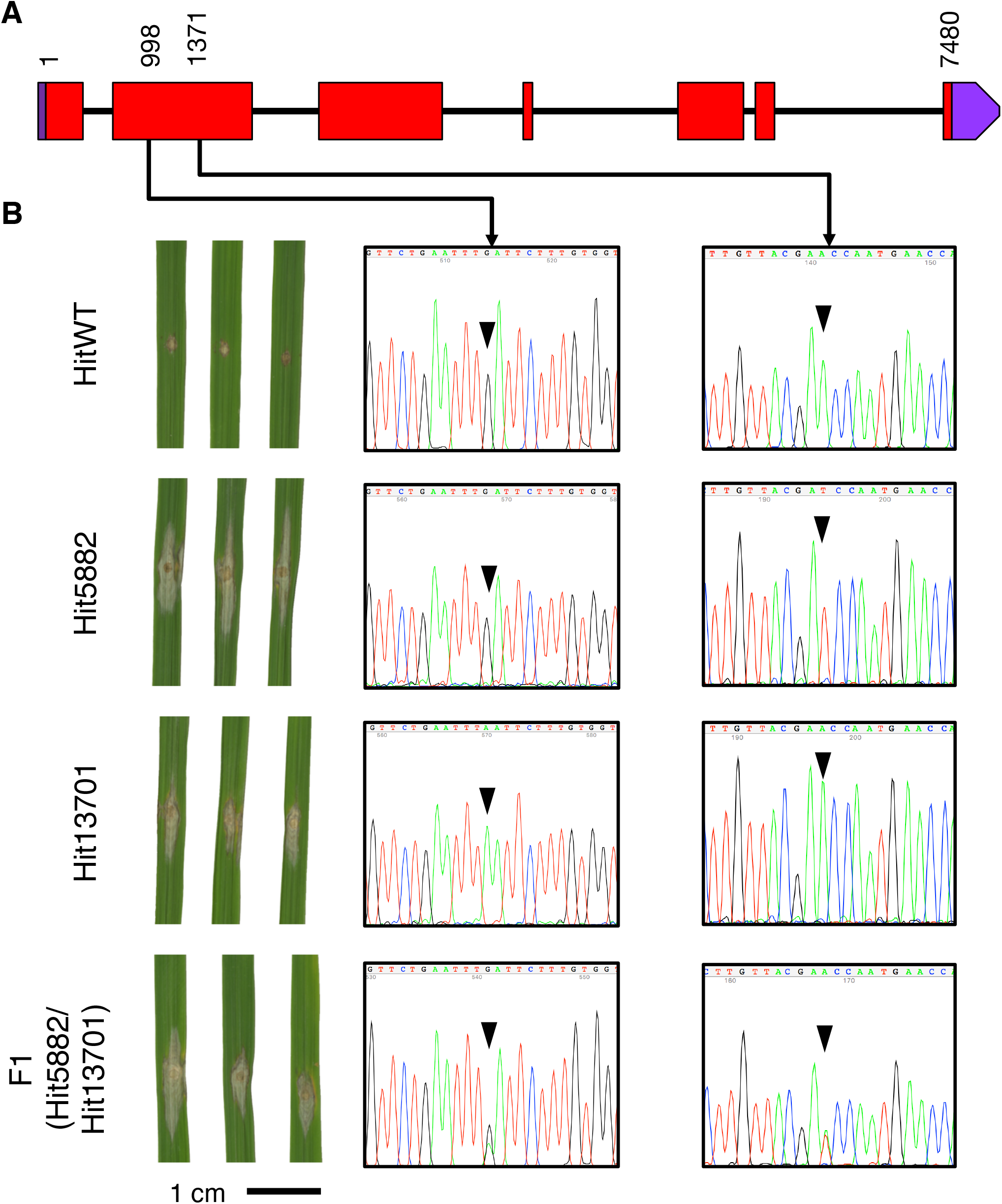
Identification of causative mutations for Hit5882 and Hit13701 by MutMap-Gap. (A) Gene structure of *Pii-2* and the SNP positions identified in Hit5882 (1371) and Hit13701 (998). Purple and red box indicates the untranslated region (UTR) and exon, respectively. Introns are indicated by horizontal bar. (B) Allelism test for Hit5882 and Hit13701 mutations. The F1 plants obtained from a cross between Hit5882 and Hit13701 were subjected to the punch inoculation with *M. oryzae* isolate TH68-126 carrying *AVR-Pii*. Heterozygosity of the two mutations of Hit5882 and Hit13701 in the F1 plants were confirmed by Sanger sequence. Arrowheads indicated the candidate mutations for Hit5882 and Hit13701. All F1 plants (Hit5882/Hit13701) showed susceptibility, suggesting that the mutations in the newly identified *NLR* gene is responsible for the Hit5882 and Hit13701 phenotypes.

To verify whether the disruption of this NLR gene in the contig “no.298” is responsible for the loss of Pii-mediated resistance in the two independent Hitomebore mutant lines, Hit5882 and Hit13701, we carried out an allelism test using F1 plants derived from a cross of Hit5882 and Hit13701. When the F1 plants derived from Hit5882 × Hit13701 were challenged with *M. oryzae isolate* TH68-126 with *AVR-Pii*, all three F1 plants showed the susceptible phenotype (Fig. 3). This result strongly suggests that the *NLR* gene in contig no.298 is required for the Pii-mediated resistance in Hitomebore. Since this *NLR* gene is located close to the previously-identified *Pii* gene (Fig. 2; Takagi *et* al., 2013), we named the newly identified gene as *Pii-2*. After confirming the sequence by Sanger sequencing, we registered this new NLR gene in NCBI (accession no LC190730) (Fig. S5). Accordingly, the gene defined as *Pii* in the previous study (Takagi *et al*., 2013) was renamed as *Pii-1* (accession no AB820896).

The *Pii-2* mutations in Hit5882 and Hit13701 are located at nucleotide-binding site (NBS) (Fig. S5). Complete loss of function of *Pii-2* is expected by the nonsense mutation in Hit13701. On the other hand, the mutation in Hit5882 causes an amino acid substitution (V371D). Although this position is distant from the P-loop of NBS required for NLR function (Mestre and Baulcombe, 2006), the drastic effect of V371D mutation (Figs. 1 and S1) indicates the importance of V371 for Pii-2 activity.

A BLAST search revealed that *Pii-1* and *Pii-2* in Hitomebore showed a high amino acid sequence similarity with *Pi5-1* and *Pi5-2* (Lee *et al*., 2009), respectively, among the known rice *NLR* genes (Fig. S6), indicating that *Pii* is allelic to *Pi5*. Between Pii and Pi5, we see some amino acid differences (Fig. S6). Currently, we do not know their importance in recognition specificity to different *AVR* alleles. The ratio of nonsynonymous substitutions to synonymous substitutions (dN/dS) between *Pii-1* vs *Pi5-1 and Pii-2* vs *Pi5-2* were 2.87 and 2.34, respectively. Higher levels of nonsynonymous substitutions as compared to synonymous substitutions implies that diversion between Pii and Pi5 may have involved positive selection.

Successful isolation of *Pii-1* and *Pii-2* by MutMap-Gap has demonstrated that MutMap-Gap enables identifying the genes that are absent from publicly available reference genome sequences (*e.g*. Nipponbare reference genome in rice). Genome organization around the rapidly-evolving genes including *NLRs* are known to be highly variable (Yang *et al*., 2013). Therefore, MutMap-Gap may be especially useful to identify *NLR* genes required for plant immunity. In this study, local *de novo* assembly for MutMap-Gap was carried out by DISCOVAR *de novo* using paired end sequence reads with 250 bp overlaps. This local *de novo* assembly achieved complete assembly of *Pii-1* gene in the contig “no.51” while *Pii-2* sequence was partial. Recent advance in PacBio sequencer (Au *et al*., 2012) will permit to generate much longer sequence reads than illumina sequencer, which may allow improvement of local *de novo* assembly. In future study we need to address the amount and length of sequence reads needed to perform high accuracy local *de novo* assembly by MutMap-Gap to apply to other crops with lager genome sizes.

In this study, we showed genetic evidence that not only *Pii-1* but also *Pii-2* is an essential component in Pii-mediated resistance. Resulting *Pii-2* deficient mutants (Hit5882 and Hit13701) together with *Pii-1* deficient mutants (Hit5948 and Hit6780) will enable us to perform experiments to analyze functions of *Pii-1* and *Pii-2*.

## EXPERIMENTAL PROCEDURES

### Mutant screening

A total 5,600 Hitomebore EMS mutant lines were applied to spray inoculation test with the incompatible rice blast (M. *oryzae*; isolate TH68-126) carrying *AVR-Pii*, as described in Takagi *et al*. (2013). The mutants screened by spray inoculation test were confirmed their phenotype by paunch inoculation test using the compatible isolate Sasa2 (*AVR-Pii*-) and the incompatible isolate Sasa2-AVRPii (AVR-Pii+) which is generated by transformation of Sasa2 with *AVR-Pii* (Yoshida *et al*., 2009).

### MutMap-Gap analysis for identifying candidate gene

MutMap-Gap analysis was carried out as described in Takagi *et al*. (2013). First, the genomic region harboring a candidate mutation was identified by the conventional MutMap method (Abe et al., 2012). Namely, the identified mutants were crossed with HitWT to generate the F2 segregating population. The F2 progeny were tested for their phenotypes by punch inoculation with an incompatible *M. oryzae* isolate. The DNAs obtained from the susceptible F2 individuals were bulked in equal amount and applied to library construction for illumina sequencer using TruSeq DNA PCR-Free LT Sample Prep Kit (illumina). The libraries were subjected to whole genome sequence by illumina sequencer NextSeq500. The MutMap analysis is carried out by pipeline version 1.4.4 (<u>http://genome-e.ibrc.or.jp/home/bioinformatics-team/mutmap</u>). In this MutMap analysis, the SNP positions where HitWT sequence shows low read depth (<=5) were excluded. Following MutMap analysis, local *de novo* assembly was carried out to reconstruct Hitomebore specific genome corresponding to the candidate genomic region. For local *de novo* assembly, the sequence reads mapped to the candidate region and those unmapped were combined and subjected to assemble by DISCOVAR *de novo* (Weisenfeld *et al*., 2014). Finally, the sequence reads from the bulked DNA obtained from the F2 progeny showing mutant phenotype were aligned to the assembled contigs and SNP-index was calculated by MutMap pipeline. A flow chart of the procedure is given in Fig. S2.

To determine the full-length coding sequence of *Pii-2* gene, the 5’- and 3’-terminal sequences of *Pii-2* cDNA of the rice cultivar Hitomebore were determined by SMARTer^®^ RACE cDNA Amplification Kit (Takara) using the gene specific primers 5’-GCAGACCTTTCCGGAGTGCTGATTGTCC-3’ (for 5’-RACE) and 5’-AACAGCAGCCTCACTGAAATTGCGAAGC-3’ (for 3’-RACE). Based on the sequence information of the 5’- and 3’-UTRs of *Pii-2* cDNA, a cDNA fragment containing the full-length coding region of *Pii-2* gene was amplified by RT-PCR using a primer set 5’-ATCCAGCGATA CAATACCA GAAGT-3’ and 5’-GCTGTGCCCAACCCCCATTTCAAT-3’. Resulting cDNA was cloned using Zero Blunt TOPO PCR cloning kit (Thermo Fisher Scientific) and the DNA sequence was determined by Sanger sequencing.

To determine the internal sequence of *Pii-2* gene between contig 298 and contig 1164 (corresponding to the 2,582-3,255 bp region of *Pii-2* cDNA coding sequence), PCR was carried out using genomic DNA as template and the primers 5’-CGATGGCTTCTCAGGTTGG-3’ (corresponding to the internal region of *Pii-2* cDNA) and 5’-TCCGGCACTGGTTTTGTG-3’ (corresponding to contig 1164), and the resulting PCR product was subjected to Sanger sequencing (fragment 1). Based on the obtained sequence information, additional genomic PCR using the primers, 5’-CCTCTCTATCACAAAACAAAACGAC-3’ (corresponding to contig 298 sequence) and 5’-TGGCAAACAGCAGAGCATG-3’ (corresponding to the internal genomic region of *Pii-2*) was performed, and the resulting PCR product was sequenced (fragment 2). DNA sequences of fragments 1 and 2 were assembled and used for reconstruction of the entire genomic sequence of the *Pii-2* gene.

## ACKNOWLEDGEMENTS

We thank Dr Kentaro Yoshida (Kobe university) and members of our lab for discussions and general assistance. This work was supported by Science and Technology Research Promotion Program for Agriculture, Forestry, Fisheries and Food Industry, Japan, Grant-in-aid for MEXT (Scientific Research on Innovative Areas 23113009) and JSPS KAKENHI (Grant No. 26850029) and JSPS Kakenhi 15H05779.

